# Viscoelasticity drives EMT in pancreatic intraepithelial neoplasia

**DOI:** 10.64898/2026.02.11.705320

**Authors:** A. Pascucci, SA. Karim, JP. Morton, M. Vassalli, M. Walker

## Abstract

Pancreatic intraepithelial neoplasia (PanIN) is a precursor of pancreatic adenocarcinoma (PDAC) and therefore critical to understand for identifying early-stage diagnostic and therapeutic targets. During PanIN, epithelial-to-mesenchymal transition (EMT) of pancreatic epithelial cancer cells is a crucial event which promotes invasion and early dissemination of cells into circulation before the full development of PDAC tumours. Changes in tissue mechanics are apparent during progression from PanIN to PDAC and increased local and global elasticity has been mathematically modelled in PanIN tissue as a predictive tool for diagnostics and development of personalized therapies. Aside from elasticity, viscoelasticity is emerging as a key feature of cancer which affects tissue mechanics through a combination of elastic and viscous components. Viscoelasticity has recently been shown to drive mechanosensitive cell behaviour and is known to change dramatically in PDAC progression. Hydrogels, as water-swollen polymer networks, are effective extracellular matrix (ECM) models that can recapitulate the viscoelastic properties of natural tissue. Despite this, hydrogels developed for studying cell behaviour in PanIN use purely elastic materials or have neglected the viscous component. Here, using PDAC mouse models, we show that viscoelasticity dynamically alters between healthy and PanIN-bearing tissue and have decoupled the role of elasticity and viscosity during EMT of pancreatic epithelial cancer cells using two-dimensional (2D) polyacrylamide (PAAm) hydrogels. Our work shows viscosity is critical in driving phenotypic changes associated with EMT in a pancreatic epithelial cancer cell line. These findings identify viscosity as an integral component of cell mechanosensing as PanIN develops, which may contribute to initial metastatic events via dissemination from the developing primary tumour. This should be explored further to potentially reveal novel diagnostic and therapeutic targets.

## Introduction

Pancreatic ductal adenocarcinoma (PDAC) is one of the most lethal cancers worldwide and contributes to >450,000 deaths per year; the prognosis is extremely poor with only 9% of patients reaching 5-year survival which is largely attributable to late diagnosis and high chemotherapeutic resistance (1-3). A significant issue with PDAC is limited early symptoms and thus lack of early-stage markers; consequently, tumours are typically only detected at advanced stages which dramatically reduces the capability for clinical intervention with only ∼15% being surgically resectable (3).

PDAC originates from normal pancreatic ductal or acinar cells that acquire mutations over time, resulting in pancreatic intraepithelial neoplasia (PanIN), which is a proposed precursor of PDAC (4). Epithelial-to-mesenchymal transition (EMT) is a critical event in PDAC, and cancer progression in general, which has been identified even in early PanIN lesions to facilitate invasion and early dissemination of pancreatic cancer cells before the full development of PDAC tumours (5). EMT is characterised as a transient, reversible phase in which epithelial cells lose their cell polarity and cell–cell junctions and develop migratory, invasive behaviours that resemble a mesenchymal phenotype which can facilitate drug resistance and metastasis (6, 7). Associated morphological and biochemical changes include changes in cell phenotype and expression patterns of key markers; for instance, cells become significantly larger during EMT and express upregulated cytoskeletal proteins such as vimentin (8). EMT is a critical process physiologically, for example, during tissue morphogenesis and wound healing. As tumours progress, it is thought that abnormal EMT can lead to intermediate quasi-mesenchymal cells with the ability to serve as cancer stem cells (9).

An indicative marker of PDAC is the development of a highly fibrotic stromal niche, known as desmoplasia, which accumulates through aberrant deposition of densely crosslinked extracellular matrix (ECM) (10, 11). Because of desmoplasia, significant changes in tissue mechanics occur; this is known to promote tumour malignancy through dysregulated intracellular signalling and higher cell contractility which drives the increased tensile state of the microenvironment (12). Several molecular players associated with cell contractility have been implicated in PDAC; for instance, focal adhesion kinase, inhibition of which increases PDAC receptiveness to immunotherapy (13). Increased local and global elasticity has been attributed to PanIN and mathematically modelled as a predictive tool for diagnostics and development of personalized therapies (14). In addition to tissue elasticity, alterations in viscous mechanical properties have been measured in PDAC and pancreatitis, which can lead to PanIN formation in the presence of aberrantly activated KRAS (15). Collectively, the elastic and viscous nature of tissues confers viscoelasticity. Viscoelastic tissue mechanics have been shown to regulate cell mechanobiology during cancer and can be effectively modelled using hydrogels (16-18). The role of viscosity in regulating EMT during PanIN progression remains to be elucidated. Given the key role of viscoelasticity in cancer progression, its investigation in PanIN could reveal novel mechanistic insights into the mechanobiology of PDAC progression that could inform therapeutic and diagnostic discovery in the future.

In this work, we demonstrate the development of several polyacrylamide (PAAm) hydrogels as two-dimensional (2D) surfaces with controlled viscoelastic properties. Using these gels, we show that the viscous contribution of the ECM is a significant driver of phenotypic changes associated with EMT in pancreatic epithelial cancer cells.

## Results

### PanIN tissue displays altered viscoelastic mechanics compared with healthy pancreatic tissue from mouse models

To characterise the changes in viscoelastic mechanical properties during the transition from healthy to PanIN tissue, we used normal healthy pancreatic tissue from C (*Pdx1-Cre*) mice (19) and PanIN-bearing tissue from KC PanIN mice (*Pdx1-Cre*; *LSL-Kras*^G12D/+^) (20). The *Pdx1-Cre* allele facilitates the recombination of target genes throughout the pancreas, as the PDX1 transcription factor is expressed in pancreatic progenitor cells from embryonic day 8.5 (21). Pancreatic cancer phenotypes are typically induced through the introduction of activating mutations in the *Kras* oncogene, and can be accelerated by deletion or mutation of tumour suppressor genes, for example, *Trp53* (22). These mouse models are highly effective at recapitulating human pancreatic cancer since *TP53* and *KRAS* mutations are present in 75% and 95% of cases respectively (23).

Atomic force microscopy (AFM) was used to assess the viscoelastic mechanical properties of the tissue samples using microindentation and microrheology measurements to derive elastic and viscous properties respectively. The workflow for sample preparation and conceptualisation of AFM measurements is shown in **Figure 1a**. We observed a homogenous distribution of stiffness values for healthy tissue samples at ∼100 Pa. In PanIN-bearing tissue, the stiffness values were more heterogeneous, with a broader distribution and areas with higher values than those seen in healthy pancreas (**Figure 1b (i)**). A higher tan δ, which indicates a greater degree of viscosity relative to elasticity, was also observed for PanIN tissue compared with healthy tissue (**Figure 1b (ii)**).

**Figure 1.**
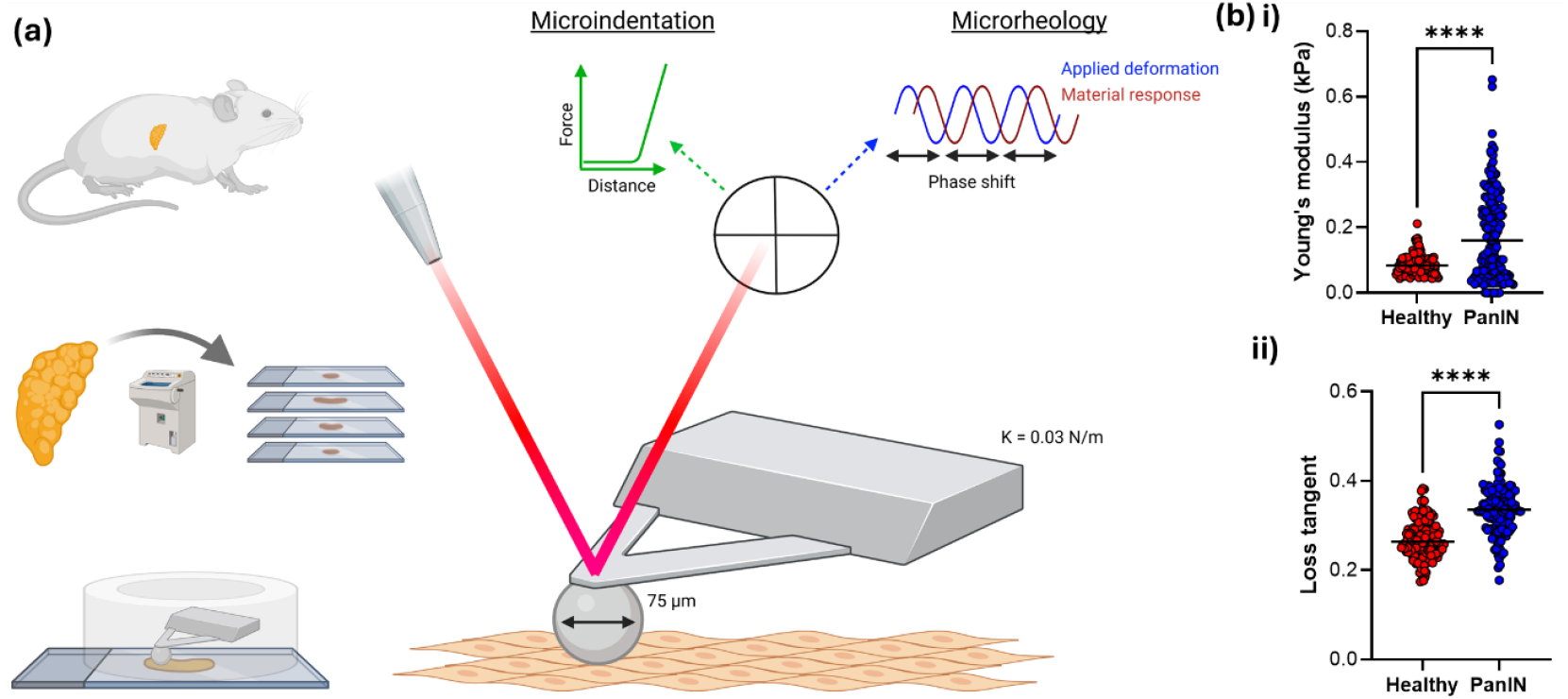
PanIN tissue displays altered viscoelastic mechanics compared with healthy pancreatic tissue from mouse models: **(a)** Schematic demonstrating the extraction and preparation process of mouse pancreatic tissue sections followed by AFM measurements of elasticity (via microindentation) and tan δ (via microrheology). **(b)** Quantification of **i)** elasticity, by microindentation, and **ii)** tan δ, by microrheology in healthy and PanIN tissue sections via AFM; n = 144-160 in which n represents each datapoint shown by circles in the graphs that correspond to an individual measurement at a position across the tissue. Individual measurements were sampled from three 100 µm^2^ areas per tissue section, with 25 points taken per area, across 3 different mice per condition. **** = p ≤ 0.0001 derived from Mann-Whitney (**i**) and unpaired t-tests (**ii**).

### Development of isoelastic PAAm hydrogels mimicking healthy and PanIN tissue elasticity shifts with altered viscous character

Having observed significant differences in healthy vs. PanIN tissue viscoelasticity, we aimed to develop PAAm hydrogels as an effective 2D culture platform with tuneable mechanics to study the effects of viscoelasticity on the mechanobiology of pancreatic epithelial cancer cells *ex vivo*. PAAm has been used as an effective tool for mechanobiology studies in PDAC and a broad range of cancers; viscoelasticity can be manipulated by altering polymer:crosslinker ratios, making it an effective material system for this study (6, 7, 18).

To study the role of viscoelasticity on pancreatic epithelial cancer cell mechanosensing and EMT, we developed a set of isoelastic gels with variable viscous character to decouple the role of viscosity from elasticity. Our strategy to modify viscoelastic mechanics of PAAm gels was to alter the ratio between polymer and crosslinker content. This approach encourages different degrees of physical entanglement within the gel network to permit variable dynamic and dissipative processes in response to mechanical stress (**Figure 2a (i)**). PAAm hydrogels were developed to mimic shifts in elasticity observed in tissue samples measured in **Figure 1** and were manipulated to establish more elastic (E) and more viscous (V) counterparts (**Figure 2a (ii)**); the specific ratios of polymer: crosslinker used to formulate each gel can be found in the **Materials and Methods** section. Within their respective (E) and (V) groups, we engineered hydrogels that retained softer mechanics to represent more healthy tissue stiffness, named “healthy”, and stiffer mechanics to mimic more PanIN-bearing tissue named “PanIN” (**Figure 2b (i)**). Within the respective (E) and (V) groups, we could also decouple the viscous from the elastic contribution of the gels shown by a significant change in tan δ but no differences in elasticity. (**Figure 2b (ii)**). These models were taken forward to explore the role of elasticity and viscosity, independently, on pancreatic epithelial cancer cell mechanosensing and EMT responses. PAAm hydrogels were also modified with 2 mM of Arginylglycylaspartic acid (RGD) peptide to facilitate cell adhesion.

**Figure 2.**
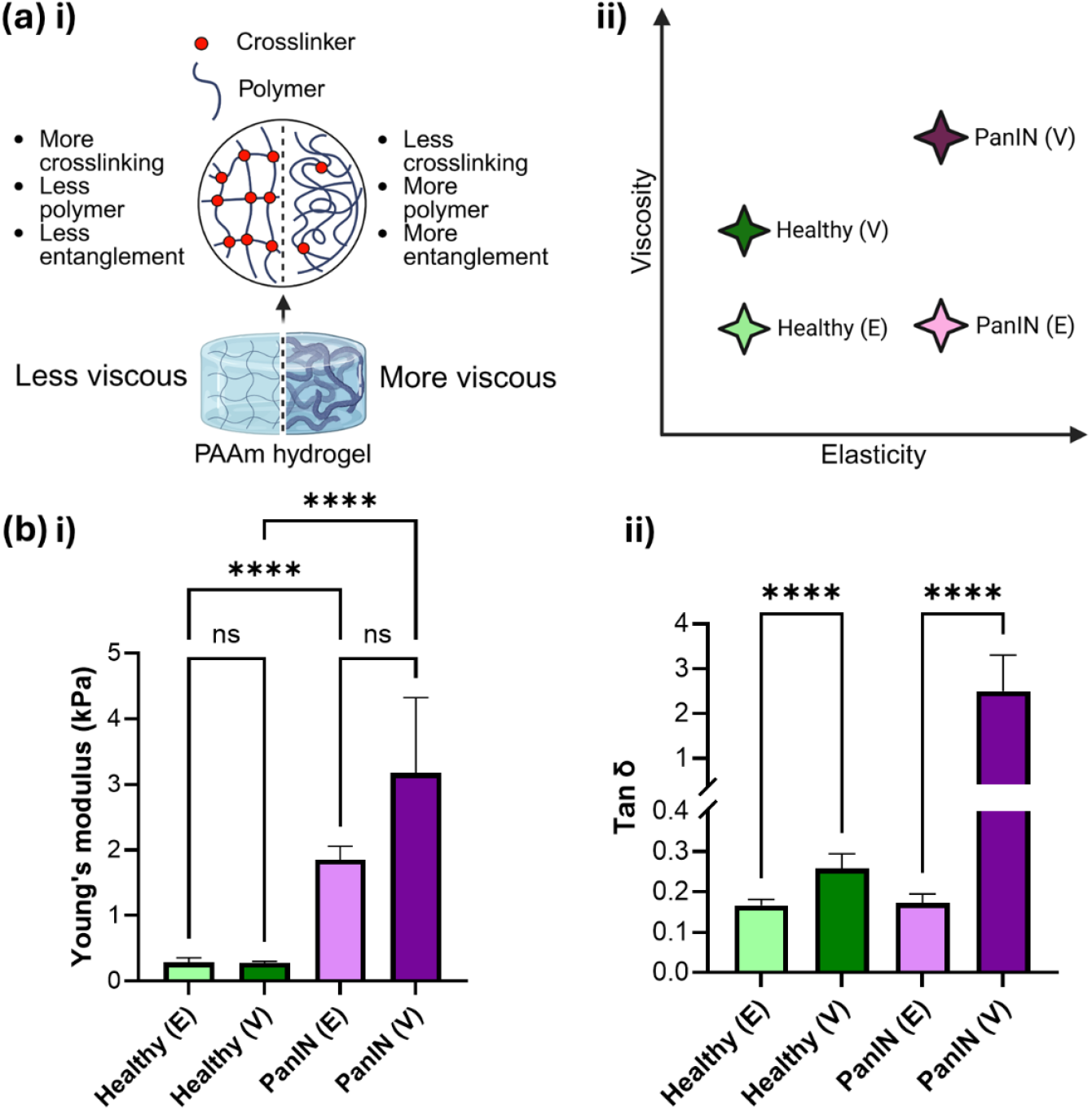
Development of isoelastic PAAm hydrogels mimicking healthy and PanIN tissue elasticity shifts with altered viscous character: **(a)** Strategy to modulate PAAm hydrogel viscoelasticity and conceptual influence on mechanics; **i)** Schematic showing the strategy for manipulating PAAm hydrogel viscoelasticity by modulating ratios of polymer:crosslinker to establish different degrees of physically entangled polymer chains, **ii)** representation of mechanics within healthy and PanIN (E/V) PAAm gel counterparts. **(b)** AFM mechanical characterisation of PAAm hydrogels with tuneable elastic and viscous properties; **i)** elasticity quantification via microindentation, **ii)** tan δ quantification via microrheology. Across all figures, n = 48-72 in which n represents individual measurements sampled from three 35 µm^2^ areas per hydrogel, with 25 points taken per area, across 3 different hydrogels per condition. Error bars denote standard deviation. **** = p ≤ 0.0001 derived from Kruskal-Wallis tests.

### Pancreatic epithelial cancer cells display morphological changes associated with EMT on stiffer hydrogels with greater viscosity

Following our mechanical characterisation of the developed PAAm hydrogels, we seeded pancreatic epithelial cancer cells on these surfaces and characterised their morphological behaviour over a series of timepoints. Across the hydrogels, we observed minimal changes at day 0 followed by some slight elongation after 3 days of culture on the PanIN (V) hydrogels; after 7 days, there were significant increases in cell size, elongation and contractility on the PanIN (V) hydrogels which are consistent with an EMT phenotype (**Figure 3a**). Intriguingly, after 7 days on PanIN gel models, we observed that the increased viscosity of the environment was the key contributor to facilitating these morphological changes; increasing the elasticity alone was insufficient to elicit a significant change (**Figure 3b (i)**). Upon quantification of morphological shape descriptors at 7 days, we observed a significant increase in cell area and concomitant decrease in cell circularity, over time for the PanIN (V) condition (**Figure 3b (ii)**). Collectively, we only observed a morphological phenotype associated with mesenchymal cells on stiffer hydrogels with greater viscosity indicating a key role of the viscous contribution in facilitating EMT within PanIN-bearing tissue (**Figure 3c**).

**Figure 3.**
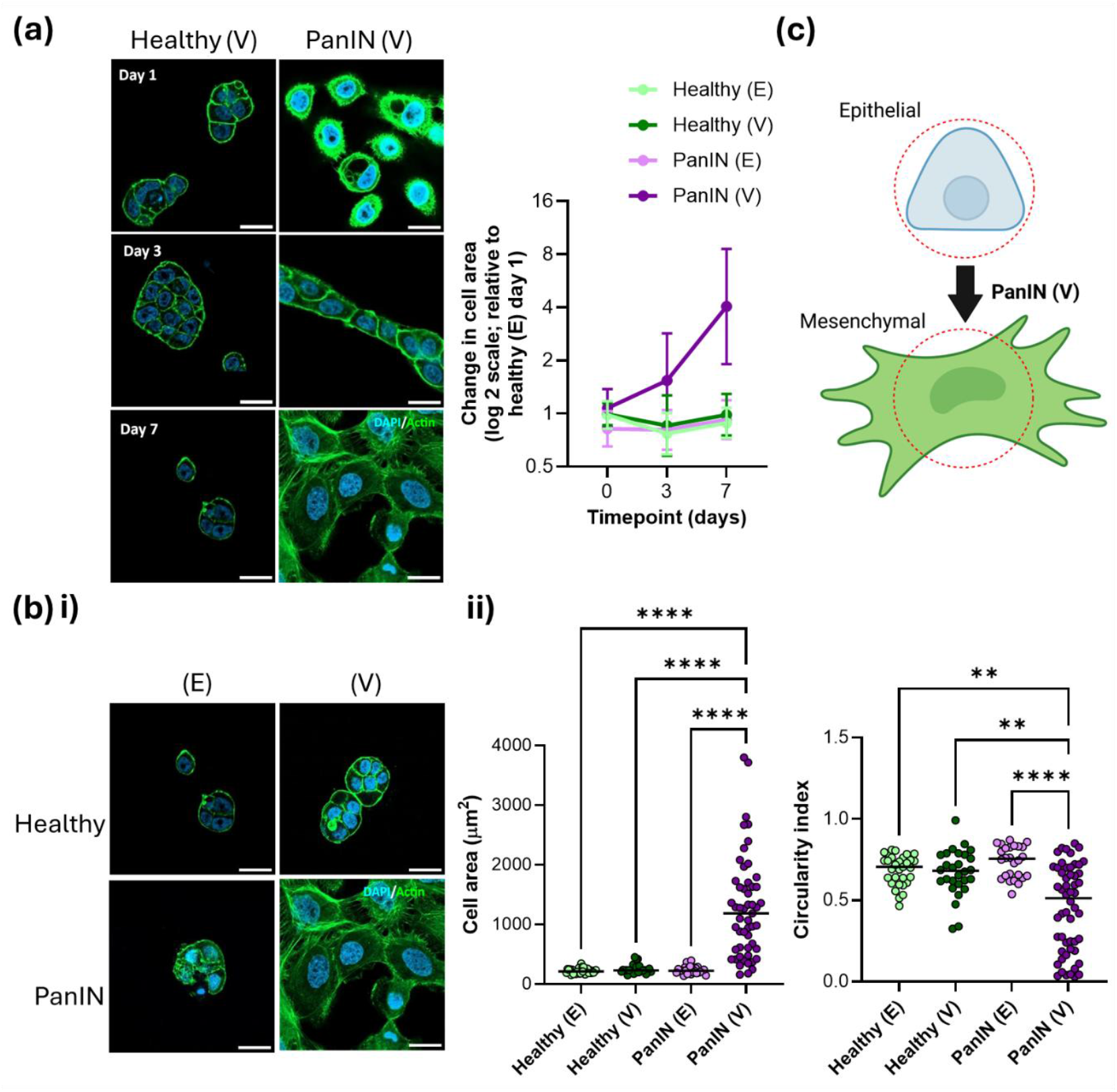
Pancreatic epithelial cancer cells display morphological changes associated with EMT on more viscous hydrogels models of PanIN-bearing tissue: **(a)** Representative images of immunofluorescent staining for actin (green) and nuclei stained with DAPI (blue), from pancreatic epithelial cancer cells grown on hydrogels over 7 days for (V) samples (left), and quantification of cell area on all hydrogels across 7 days (right). **(b)** Cell morphology on hydrogels at 7 days; representative images of immunofluorescent staining for actin (green), and nuclei stained with DAPI (blue), from pancreatic epithelial cancer cells grown on all hydrogels at 7 days (**i**) with quantification of cell area (left) and circularity (right) (**ii**). **(c)** Schematic representation of morphological descriptors as a readout of EMT. Across all figures, n = 26-54 in which n represents individual cell measurements sampled from random areas on each hydrogel across three technical replicate samples per condition, error bars denote standard deviation, ** = p ≤ 0.01, **** = p ≤ 0.0001 derived from Kruskal-Wallis tests. Scale bars = 30 µm.

### Pancreatic epithelial cancer cells upregulate cytoskeletal and mechanotransductive EMT markers on stiffer hydrogels with greater viscosity

Following our observations of altered pancreatic epithelial cancer cell morphology in response to PAAm hydrogels with altered viscoelasticity, we explored the relationship with known mechanosensitive proteins associated with EMT. Yes-associated protein 1 (YAP) has been described as a mechanical rheostat which is heavily implicated in EMT during abnormal cell mechanosensing in PDAC and cancer generally (6, 7, 24). We observed significantly increased nuclear translocation of YAP in pancreatic epithelial cancer cells grown on PanIN (V) substrates compared to softer or more elastic conditions (**Figure 4a**). Additionally, the intermediate filament protein vimentin is a characteristic EMT marker that takes an active role as a mechanosensor during metastasis by interacting with other proteins to provide dynamic viscoelastic support to the cell (25). As with YAP, we observed a significant upregulation of vimentin expression in pancreatic epithelial cancer cells grown on PanIN (V) substrates compared with healthy and more elastic conditions. We also observed a significant increase in vimentin expression in cells on healthy (V) or PanIN (E) substrates compared to healthy (E) substrates suggesting a role for increased elasticity, or softer tissue viscosity, in upregulating vimentin expression levels (**Figure 4b**). Collectively, we have identified a critical role for viscoelasticity within the PanIN microenvironment to drive protein expression changes associated with EMT which facilitate loss of cell-cell polarity and increased invasive potential (**Figure 4c**).

**Figure 4.**
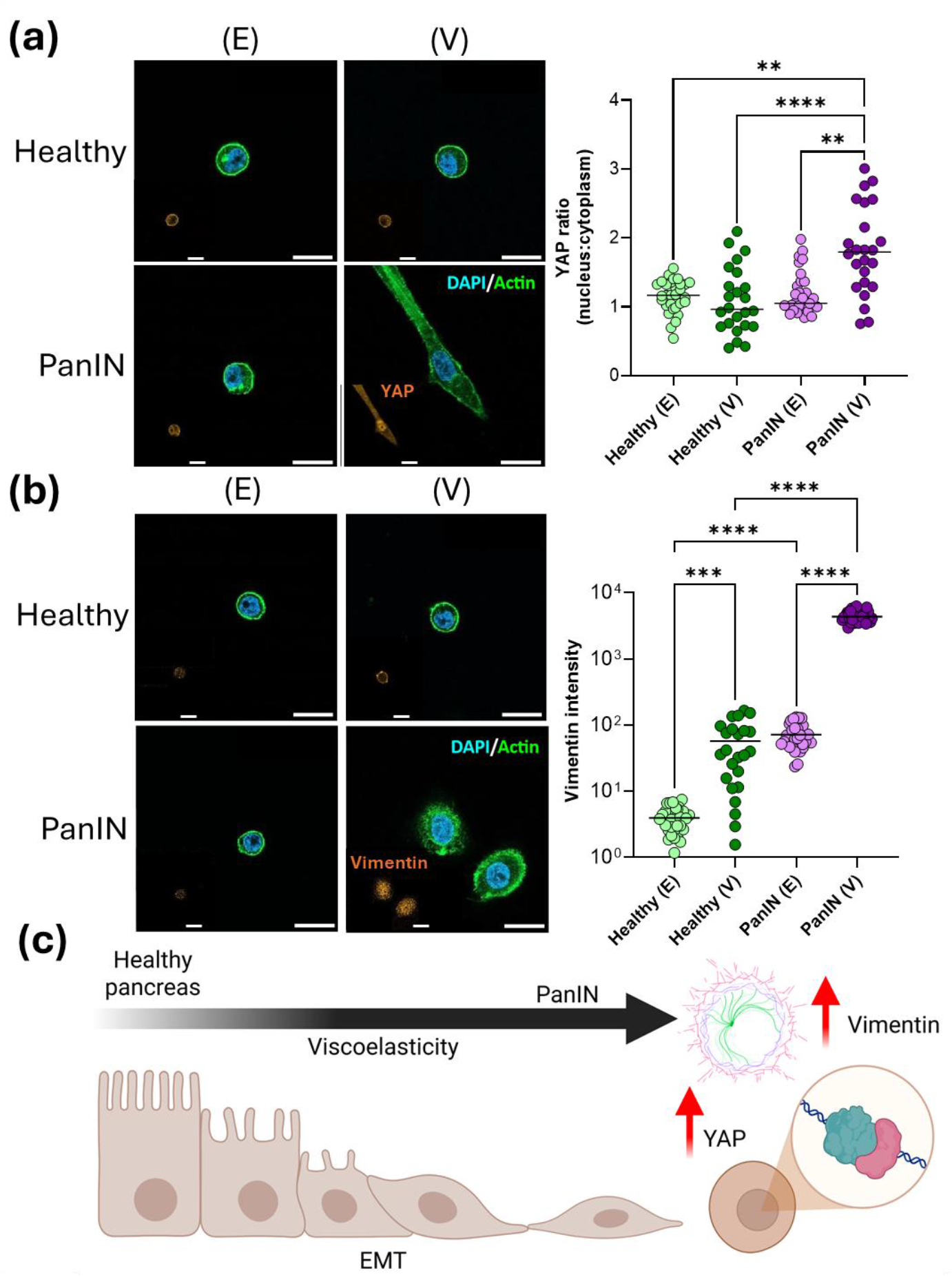
Pancreatic epithelial cancer cells upregulate cytoskeletal and mechanotransductive EMT markers on stiffer hydrogels with greater viscosity: **(a)** Characterisation of YAP localisation within pancreatic epithelial cancer cells after 3 days culture in response to altered hydrogel viscoelasticity. Representative images of immunostaining of cells with DAPI, or antibodies against actin (green) and YAP (red, left) and quantification of nuclear:cytoplasmic YAP ratios (right). **(b)** Characterisation of vimentin expression within pancreatic epithelial cancer cells after 3 days culture in response to altered hydrogel viscoelasticity. Representative images of cell immunostaining with DAPI, or antibodies against actin (green) and vimentin (red, left) and quantification of vimentin intensity normalised to healthy (E) sample (right). **(c)** Schematic to represent the role of viscoelastic cell mechanosensing during EMT in PanIN tissue through upregulation of YAP and vimentin. Across all figures, n = 24-33 in which n represents individual cell measurements sampled from random areas on each hydrogel across three technical replicate samples per condition, ** = p ≤ 0.01, *** = p ≤ 0.001 **** = p ≤ 0.0001 derived from Kruskal-Wallis (**i**) and Brown-Forsythe and Welch ANOVA tests (**ii**). Scale bars = 30 µm.

## Discussion

In this study, we decouple the roles of viscosity and elasticity on EMT during the PanIN stage, that precedes PDAC, by characterising the viscoelastic properties of healthy and PanIN tissue. We assessed this by developing PAAm hydrogels to recapitulate changes in viscoelasticity during the transition from healthy to PanIN-bearing tissue and assessing pancreatic epithelial cancer cell EMT in response to altered substrate mechanics. We highlight the critical role that viscosity specifically has on EMT during the complex viscoelastic changes that occur in tissue mechanics during PDAC progression. These findings begin to address the intricate nature of cell mechanosensing during early stages of PDAC and identify viscosity as integral component of tumour progression which should be explored further to potentially reveal novel diagnostic and therapeutic targets.

Even in early PanIN lesions, EMT markers are expressed which promote invasion and early dissemination of pancreatic cancer cells before the full development of PDAC tumours, which highlights the pathological relevance of EMT at this stage (5). The inflammatory nature of damage within the pancreas facilitates physical and chemical changes in the tissue microenvironment which allows cells in early lesions to lose their ductal phenotype, detach from the epithelium, and invade the surrounding tissue (26). Understanding how viscoelasticity plays a role in this response could be critical for early detection and intervention in PDAC to ensure that initial dissemination from the developing primary tumour can be identified for therapeutic intervention. Targeting mechanosensitive cellular processes that occur in response to altered viscoelasticity may be a viable strategy to combat EMT-mediated dissemination.

Alterations in viscoelastic mechanics during the transition from healthy tissue to PDAC are well known, with higher viscous properties being present in PDAC (15). We should stress that the changes in viscoelastic mechanics that were measured in PanIN samples were organised spatially across the tissue in a heterogenous manner, which points to a highly localised mechanosensitive response and suggests precise mechanical characterisation may be required for accurate diagnostic readouts.

Our engineered PAAm hydrogels were highly effective substrates for manipulating changes in viscoelasticity that we observed in our pancreatic tissue measurements to elucidate the role of viscosity in promoting mechanosensitive EMT independently of elasticity alone. EMT is characterised by a variety of changes in protein marker expression and alterations in cell morphology. In this short study, we have uncovered several fundamental phenotypic EMT changes in response to altered matrix viscoelasticity. Future work in this space should characterise a broader range of EMT markers, as well as expected changes in behavioural cell response, such as migration and invasion. We observed gradual phenotypic changes over 7 days which confirms the mechanosensitive plasticity of pancreatic epithelial cancer cells under these in vitro conditions.

We highlight how matrix viscoelasticity promotes different aspects of EMT, as shown in **Figure 4c**, which has great importance for developing our understanding of PDAC progression. The development of therapeutics that perturb mechanosensing of cells in response to viscoelastic changes within the tumour at an early stage will benefit from prior understanding of the nature of the specific cellular responses that occur during this phenomenon. It is critical that further studies are carried out to establish whether altered matrix viscoelasticity facilitates altered cell responses in synergy with other cues within the tumour microenvironment. For instance, after more thorough 2D characterisation, the dimensionality of the models should eventually be transferred to more complex three-dimensional (3D) systems. 3D systems could benefit from label-free techniques that our group have previously established to mechanically phenotype cell spheroids in 3D hydrogel models; these may present diagnostic opportunities for characterising micrometastases within PanIN-bearing tissue models (27). Additionally, co-cultures with other relevant native cell types should be included, such as pancreatic stellate cells, which significantly contribute to cell-cell signalling and remodelling of the local stroma to develop a fibrotic desmoplastic niche. These adaptations will more accurately recapitulate the spatiotemporal and cellular environment of pathological in vivo tissue (28, 29). Adding complexity when modelling tissue at early stages of PDAC progression will also give more insight into chemotherapeutic drug resistance. For instance, previous work using 2D models has shown that resistance profiles to gemcitabine and paclitaxel vary during mechanosensitive EMT, which could be affected by other factors in vivo that further induce EMT towards a more stable mesenchymal phenotype (30).

## Materials & methods

### Genetically engineered mouse models (GEMM)

*Pdx1-Cre*; *Kras*^+/+^ mice and *Pdx1-Cre; LSL-Kras*^G12D/+^ mice were bred in house at the CRUK Scotland Institute on a mixed background. Animals were maintained in conventional caging with environmental enrichment, continual access to standard irradiated diet and water, and on a 12-hour light dark cycle. Mice of both sexes, aged between 6 and 18 weeks old, were used in approximately equal numbers. Genotyping was performed by Transnetyx (Cordoba, TN, USA). Animals were euthanised at timepoints for tissue harvest and pancreata were collected and snap frozen. All studies were conducted under UK Home Office licence and approved by the University of Glasgow Animal Welfare and Ethical Review Board (AWERB).

### Histology

Optimal Cutting Temperature (OCT)-embedded samples were sectioned with a cryostat as fresh 10 μm sections; fixation was not performed to preserve native mechanical properties.

### Atomic force microscopy

Using a NanoWizard 3 Bioscience AFM (JPK, Berlin, Germany), calibrations were conducted in specific aqueous medium for each type of sample; water was used for PAAm gels and protease inhibitor cocktail solution (Merck) diluted 1:100 in phosphate buffered saline (PBS) was used for tissue samples to prevent proteolytic tissue degradation. Measurements and calibration of cantilever sensitivity against a stiff surface (tissue culture dish) and of spring constant, using the thermal noise method, were done using the JPK SPM software (version 6.1.192).

Force spectroscopy measurements were performed using a constant cantilever approach speed of 5.0 µm/s. 0.03 N/m ARROW-TL1 cantilevers (NanoWorld) mounted with a 75 µm diameter spherical silica tip were used. Microindentation measurements were done using an indentation depth of ∼3 µm. Microrheology measurements included a pause segment of 0.5 s at constant height after a ∼3 µm indentation followed by a 0.4 s oscillation at a frequency of 10 Hz and amplitude of 10 nm; this was used to derive the viscoelastic response of the samples. From microindentation and microrheology measurements, Young’s moduli and tan δ were calculated using the Hertz model (31) and microrheology processing functions of the JPK DP software respectively (version 6.1.192).

For tissue samples, individual measurements were sampled from three 100 µm^2^ areas per tissue section with 25 points taken per area across 3 different mice per condition. For hydrogel samples, individual measurements were sampled from three 35 µm^2^ areas per hydrogel with 25 points taken per area across 3 different hydrogels per condition. Measurements were acquired from randomly selected positions across all samples.

### Glass coverslip preparation for PAAm gels

12 mm glass coverslips and cover glass slides were Radio Corporation of America (RCA) cleaned by washing in water and ethanol before heating for 10 min at 65 °C in a 5:1:1 solution of water:H_2_O_2_:NH_3_. After drying, cover glass slides were covered with Rain-X for 5 s, washed in ethanol, and dried to use as hydrophobic glass slides. RCA-cleaned 12 mm coverslips were acryl-silanized by submerging for 2 h in a 0.5 % solution of 3-(acryloyloxy)propyltrimethoxysilane (Alfa Aesar) in ethanol with 5 % water. Coverslips were then dried and tempered by incubating for 1 h at 120 °C

### PAAm hydrogel preparation

All reagents were acquired from Sigma Aldrich. Briefly, 1 mL volumes were prepared using stock solutions of 40 % acrylamide and 2 % N,N’-methylenebisacrylamide mixed in different ratios for specific gel compositions (**Table 1**). Solution volumes were then made up to 1 mL with milli-Q water, 2.5 µL tetramethylethylenediamine, and 7.5 µL 10 % ammonium persulfate and mixed thoroughly. 10 µL of solution was spotted onto hydrophobic glass slides before placing acrylsilanized glass coverslips onto the spots. Gelation was allowed to occur at room temperature for 30 min before detaching and swelling in water overnight at 4 °C.

**Table 1.** Acrylamide and N,N’-methylenebisacrylamide ratios for PAAm hydrogels.

| **Sample** | **Acrylamide concentration (%)** | **N,N'-methylenebisacrylamide concentration (%)** |
| --- | --- | --- |
| Healthy (E) | 3.75 | 0.03 |
| Healthy (V) | 10 | 0.001875 |
| PanIN (E) | 9 | 0.02 |
| PanIN (V) | 25 | 0.00125 |

### Peptide functionalisation of PAAM gels

PAAm gels prepared on coverslips were transferred to multiwell plates before covering with 0.2 mg mL−1 sulfosuccinimidyl 6-(4′-azido-2′-nitrophenylamino)hexanoate (Sulfo-SANPAH) (Thermo Fisher) solution. Samples were placed in a 365 nm ultraviolet (UV) light source at a distance of ≈3 inches and exposed for 10 min; this process was repeated three times. Gels were then washed with 50 mM HEPES buffer (pH 8.5) three times before covering with 2 mM RGD peptide solution (prepared in same HEPES buffer) and overnight incubation at 37 °C. Gels were then washed with sterile-filtered milli-Q water to remove excess peptide. UV sterilisation for 30 mins was performed in a tissue culture hood before cell work.

### Cell culture

Pancreatic epithelial cancer cells (BxPC-3, ATCC) were derived from human PDAC tissue and originate from an epithelial cell line isolated from a biopsy specimen taken from the body of the pancreas of a 61-year-old female patient; these were authenticated by short tandem repeat profiling. Cells were grown in RPMI 1640 medium with 10 % foetal bovine serum (FBS), 2 mM L-glutamine, 1 % penicillin/streptomycin (Sigma-Aldrich) and 1 % fungizone amphotericin B (Gibco). Throughout all culturing, cells were incubated at 37 °C in a 5 % CO^2^ atmosphere. For experiments, cells were initially cultured between passages 6-8 in flasks no higher than 70-80 % confluence before trypsinising and seeding onto PAAm hydrogels at a density of 10,000 cells/cm^2^. Mycoplasma testing was routinely assessed within the laboratory by assessing for visible presence of DNA staining patterns around cells.

### Immunostaining

Samples were washed with PBS before using 4 % formaldehyde for 30 min at 4 °C. Then, PBS washes were done followed by permeabilization in 0.1 % Triton X-100 for 5 min at room temperature. Samples were then washed with PBS and blocked for 1 h in 1 % bovine serum albumin (BSA). After blocking, all primary antibody (Ab) solutions were prepared in 1 % BSA at appropriate dilutions and added to cover the samples before incubating overnight at 4 °C. After primary Ab incubations, samples were washed three times with 0.5 % Tween-20 prior to addition of secondary Ab solutions (diluted appropriately in 1% BSA) and incubation at room temperature in the dark for 1 h. Samples were then washed three times with 0.5 % Tween-20 before mounting onto glass slides using VECTASHEILD antifade mounting medium with DAPI (Vector Laboratories). Visualization was done using a Zeiss LSM900 confocal microscope (Zeiss, Germany).

The following primary antibodies and dilutions were used: YAP (Santa Cruz, sc-101199, 1:250 dilution), Vimentin (Agilent DAKO, GA630, 1:300 dilution).

The following secondary antibodies and dilutions were used: Cy™3 AffiniPure® Goat Anti-Rabbit/Rabbit Anti-Mouse (Jackson ImmunoResearch, 111-165003/315-165-003, 1:400 dilution), Alexa Fluor 488 phalloidin (Thermo Fisher, A12379, 1:400 dilution).

### Image analysis

Using ImageJ (version 1.53t), images were split into three channels to separate the different stained markers. Using the threshold function, the nucleus and actin cytoskeleton images were binarized and selected to calculate morphological parameters, such as cell area and circularity, via the measure function.

YAP levels were characterized by the nuclear/cytoplasmic expression ratio, which was obtained by measuring the integrated density of the protein and normalizing relative to cytoplasmic/nuclear area, as described in Equation [1]:

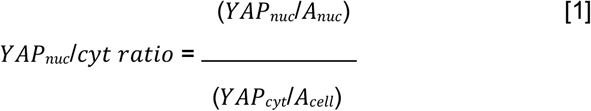

Where *YA P*_*nuc*_ and *A*_*nuc*_ are the respective expressed YAP levels in the nucleus and the nuclear area, and *YA P*_*cyt*_ and *A*_*cell*_ are the cytoplasmic YAP levels and cellular area respectively.

To quantify vimentin expression levels, the integrated density of the protein was measured and normalized to the cell area, as showed in Equation [2]:

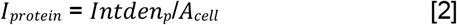

Where *I*_*protein*_ is the normalized intensity of the protein levels, *Intden*_*p*_ is the measured integrated density, and *A*_*cell*_ is the cell area.

### Statistical analysis

Using GraphPad Prism (version 10.5.0), outliers were identified using the Robust regression and OUTlier removal (ROUT) test and descriptive statistics were performed to assess whether error between samples exceeded a two-fold change to determine whether to assume equal standard deviations. The following normality and lognormality tests were realized to determine whether to run a parametric or non-parametric test: D’Agostino & Pearson, Anderson-Darling, Shapiro-Wilk, and Kolmogorov-Smirnov. Appropriate t-tests for pairwise comparisons (either unpaired t-test or Mann-Whitney) or one-way Analysis of Variance (ANOVA) tests for multiple comparisons (either ordinary ANOVA, Brown-Forsythe and Welch or Kruskal-Wallis tests) were performed with data variation being considered significant when p ≤ 0.05 (* p ≤ 0.05, ** p ≤ 0.01, *** p ≤ 0.001, **** p ≤ 0.0001).

## Acknowledgements

The authors acknowledge the Centre for the Cellular Microenvironment (CeMi, Glasgow, UK) leadership for generous allowance of general lab resources and equipment usage within the project and thank the Biological Services team at the CRUK Institute Scotland for processing and handling of mouse samples. We acknowledge funding obtained from the StemNiche grant (UKRI/EPSRC, grant number EP/X036049/1) and Scottish Universities Life Sciences Alliance (SULSA) ECR prize to support this work. SAK and JPM were supported by Cancer Research UK core funding to the CRUK Scotland Institute (A17196 and A31287) and to the Morton lab (A29996).

## Author contributions

AP performed most experimental work, data analysis and figure preparation; SAK and JPM performed GEMM work; MV provided guidance and editorial manuscript support; MW conceptualised the idea for this project, performed pancreatic tissue mechanical measurements, and wrote the manuscript with significant input from all authors.

## Data availability

All data supporting the findings reported in this paper are provided in full in the Results section of the paper

